# Dynamics of calcium oxalate monohydrate in high and low temperature phases using ^17^O solid-state NMR

**DOI:** 10.64898/2026.08.22.746468

**Authors:** Liliya Vugmeyster, Kusum Yadav, Sean T. Holmes, Dmitry Ostovsky

**Affiliations:** Department of Chemistry, University of Colorado at Denver, Denver CO 80204 USA; National High Magnetic Field Laboratory, Tallahassee, FL USA 32310 USA; Department of Chemistry & Biochemistry, Florida State University, Tallahassee, FL 32306 USA; Department of Mathematical and Statistical Sciences, University of Colorado at Denver, Denver CO 80204 USA

## Abstract

Calcium oxalates are naturally occurring minerals, with the monohydrate form CaC_2_O_4_·H_2_O (COM) being the most stable. COM contains two crystallographically different water sites. We investigate the details of water internal dynamics in the high and low temperature phases of COM using ^17^O central transition solid-state NMR line shapes, as well as laboratory and rotating frame relaxation rates. The measurements were performed either under static or magic angle spinning conditions and in a wide temperature range from 343 to 180 K. The combination of all measurements allows for precise constraints on motional mechanisms, rate constants, and amplitudes of motions. The high temperature phase is dominated by large-angle fluctuations with an amplitude of about 100*°*, identical in both sites. During the phase transition between 323 to 300 K, these large-angle jumps freeze out in one of the water sites, while remaining active in the other. In the low temperature phase from 280 to 180 K, small-angle fluctuations of 2-8*°* in amplitude dominate the relaxation. Transverse relaxation rates also point to the existence of a very slow collective “rocking” motion down to about 220-200 K.

## Introduction

Calcium oxalates are naturally occurring minerals found in very different environments, from soil and plants to sediments in lakes and oceans,[1, 2] to kidney stones.[3] The monohydrate form of calcium oxalate, CaC_2_O_4_·H_2_O (COM), is considered the most stable. Trihydrate, dihydrate, and amorphous phases undergo phase transformations into the monohydrate state.[4] The exact role of water in the phase transitions and stabilization of the monohydrate form remains a topic of active investigation. Most works have focused on structural features using computation and multiple spectroscopic techniques.[5-10] Neutron diffraction crystallography results were reported by Daudon *et al*.[10] Goldberga et al.[9] utilized ^2^H and ^17^O solid-state NMR methods to decipher the details of structures and several dynamic features of COM. We build upon their work to dive into the details of dynamics of water molecules across a wide temperature range, from 343 to 180 K, using ^17^O solid-state NMR relaxation. The ^17^O enrichment protocols for incorporation of ^17^O atoms into COM[9] are crucial to the sample preparation, enabling our dynamics studies.

COM undergoes a phase change from a low temperature form (LT) to the high temperature form (HT) at around the 328 to 348 K temperature range. The LT form[10] possesses two crystallographically different water molecules which form a dimer, denoted as W_1_ and W_2_. The main difference is that for the W_1_ molecule, two hydrogen bonds are present with oxalate, while the W_2_ molecule forms a hydrogen bond with one oxalate oxygen and with the oxygen atom of W_1_. In the HT form, the structure has been proposed to be disordered, with water molecules occupying multiple orientations (Figure 1). Four orientations were suggested based the on computational work by Shepelenko et al.[5]

**Figure 1.**
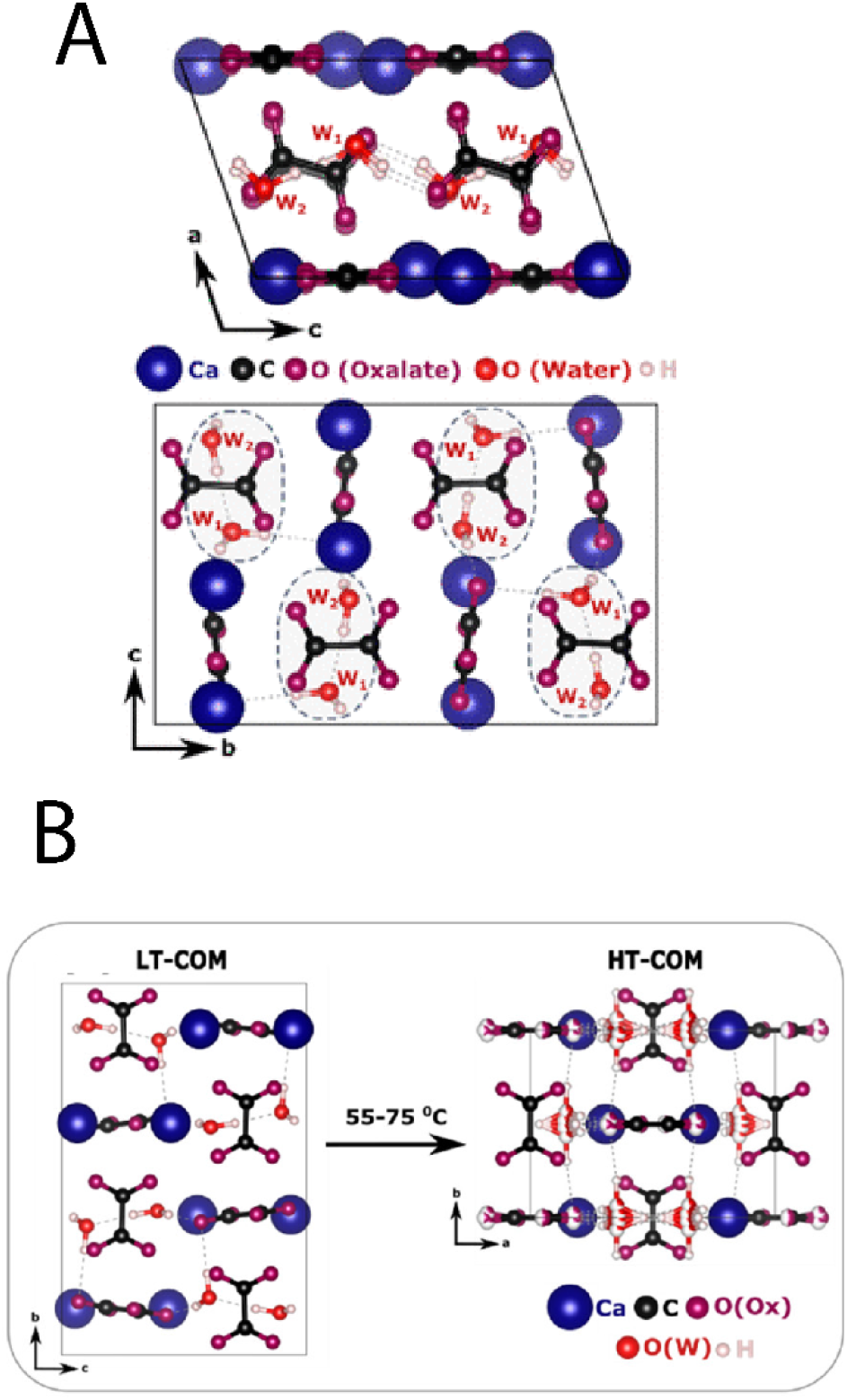
A) Crystal structure[10] of the LT phase of CaC_2_O_4_·H_2_O, with two different water sites denote by W_1_ and W_2_. B) The proposed transition from LT to HT form. Hydrogen bonding is highlighted with gray dashed lines. The 4 orientations of H-bonds suggested in[5] are shown by partial white/red coloring. Reprinted with permissions (license ID: 1762298-1) from Figures 1 and 4 of Goldberga et al.[9], CC BY-NC-ND 4.0 license, available online at https://doi.org/10.1021/acs.jpcc.2c02070.

In this work, we determine the extent of flexibility of water molecules in the LT and HT phases using ^17^O solid-state NMR relaxation and line shape analysis approaches. Goldberga et al.[9] used ^2^H solid-state NMR to probe deuterium flips in D_2_O-enriched COM and the amplitude of the twisting motions across the two phases. They also observed a significant reduction of the apparent ^17^O quadrupolar tensor coupling constant with the increase in temperature in the HT form, suggesting fluctuations of local environment around the oxygen atoms. ^17^O relaxation measurements can provide further insights into the mechanisms, amplitudes, and rate constants of the motions of water molecules. Complementary results can also be obtained from ^2^H-based studies, for which relaxation is expected to be dominated by deuterium flips in D_2_O molecules. We utilize ^17^O central transition (CT) solid-state relaxation and line shape experiments combined with computational modeling approaches developed previously[11, 12] to determine motional modes and their parameters in COM for both the HT and LT phases.

## Materials and Methods

COM with ^17^O isotopic enrichment of water molecules was provided by Prof. Danielle Laurencin, prepared as described in reference .[9] 20 mg of the sample was packed into a glass tube with 5 mm outer diameter for static measurements, and 10 mg into a 2.5 mm rotor for MAS measurements.

### NMR measurements

NMR measurements were performed at the National High Magnetic Field Laboratory (NHMFL) using a 18.8 T Bruker Avance NEO spectrometer equipped with a medium-bore Oxford magnet, and either a NHMFL-built static HX low-E probe[13] with a coil inner diameter of 5 mm or a NHMFL-built HX low-E probe with magic-angle spinning (MAS) capabilities and with a coil diameter of 2.5 mm. MAS measurements were performed at a spinning rate of 25 kHz. The temperature calibrations were performed using lead nitrate[14].[15]

The detection scheme for the static measurements was focused on the CT and utilized the Hahn echo pulse sequence,[16] with an echo time of 25 *µ*s under the static conditions, while for MAS measurements, a single pulse detection was used. The rotor-assisted population transfer (RAPT) block was employed for signal enhancement of the line shapes, utilizing the saturation of the first satellite transitions (ST1). This was achieved using selective off-resonance irradiation centered at ST1. We used a sequence consisting of ten pairs of Gaussian-shaped pulses of 10 *µ*s duration with a ±300 kHz offset, spaced at 1 *µ*s intervals. This RAPT block is an extension of the prior scheme by Haase et al.[17]. The strength of the ^17^O CT excitation and inversion pulses was 30-35 kHz. SPINAL-64 proton decoupling[18] during the acquisition period was applied with an RF amplitude of 70 kHz. ^17^O *T*_1_ measurements under static conditions were performed using the inversion recovery sequence without the pre-saturation of ST1, while ^17^O *T*_1_ measurements under MAS utilized the saturation recovery sequence due to the absence of signal inversion. *T*_2_ measurements under MAS utilized the Hahn echo scheme with the variable echo period. ^17^O *T*_1*ρ*_ measurements under static conditions utilized the pulse sequence described in references, [11, 19] with the spin-lock period of variable length and RF amplitude of 35 kHz. Continuous wave decoupling of protons with a 70 kHz RF field strength was applied during the *T*_1*ρ*_ relaxation period. 16 to 64 scans were collected, with 512 complex points and spectral window of 400 kHz. 4 to 8 dummy scans were employed in the relaxation measurements.

The spectra were processed with the 1 kHz line broadening and referenced to the natural abundance ^17^O signal of tap water at 280 K, taken as 0.0 ppm. The magnetization build-up and decay curves were constructed using the integration over the entire powder patterns. Mono-exponential *T*_1_ inversion, as well as mono-exponential *T*_1*ρ*_ and *T*_2_ magnetization decay curves, were fitted to the function *M*(*t*) = *M*_∞_ + (*M*_0_ − *M*_∞_)*e*^−*t*/*T*^, in which *M*_∞_ is an effective baseline parameter. Bi-exponential magnetization decay curves were fitted using

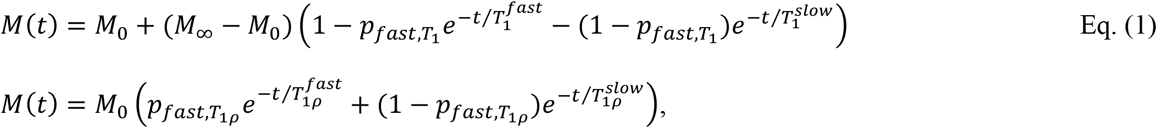

in which 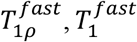 are the fast-relaxing components and 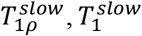 are the slow-relaxing components, and 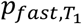and 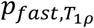 are the fractions of the components relaxing with the times 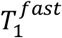and 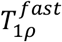 relative to the ones relaxing with the times 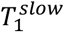 and 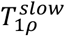 respectively.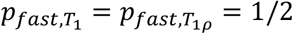 for the case of COM, reflecting equal contributions from the two water sites. For saturation recovery build up curves the same equations apply, and *M*_0_ can be an additional fitting parameter.

### Simulations

Simulations of ^17^O line shapes and relaxation curves were performed as described in prior works, [11, 12, 19-22] employing numerical integration of the Liouville – von Neumann equations by matrix exponentiation using an internal MATLAB algorithm. Relaxation delays employed in simulations matched those used in the experiments.

The following site-specific quadrupolar (EFG) and chemical shift anisotropy (CSA) tensor parameters were used, as determined by Goldberga et al.[9] using experimental line shapes under static conditions, MQMAS measurements, and DFT calculations: quadrupolar coupling constant *C*_*q*_ = 6.5 MHz for both sites; quadrupolar tensor asymmetry *η*_*q*_ = 0.83 and 0.97 for W_1_ and W_2_ sites, respectively; isotropic chemical shift *δ*_*iso*_ = 19 and 9 ppm; chemical shift anisotropy Δ*δ*_*CS*_ = *−*45 and 35 ppm; and chemical shift asymmetry *η*_*CS*_ = 0.6 and 0.7 for W_1_ and W_2_ sites, respectively. The ZCW tiling algorithm[23, 24] was used with 15,000-50,000 tiles for each site.

Three individual types of motions were set up relative to the principal axis system (PAS) frame for the ^17^O EFG tensor. CSA tensor parameters were then defined relative to the EFG PAS frame and followed it throughout. See Vold and Hoatson[20] for details on setting up motional frames. The parameters of the tensors for the two sites were assigned arbitrarily as W_1_ for a less mobile site A and W_2_ for a more mobile site B (see below). All relaxation rates change values by less than 5% if this assignment is reversed.

Large-angle jumps occur between two sites with the same orientation of one of the O*−*H bonds and rotation angle *α* around it. The orientations are given by (0,0,0) and (*φ, α*, 2*π* − *φ*), where 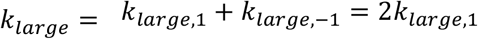. The rate constant 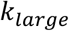 is defined as a sum of the forward and backward jump rates 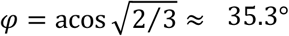, where the forward and backward rates are taken as equal.

The small-angle fluctuations were modeled as taking place around the H*−*H direction, which coincides with the *y*-axis of the PAS frame for ^17^O EFG tensor. These fluctuations are modelled with 3-site jumps between sites with equal weights. Their coordinates given by (0, *θ*, 0), (0,0,0), (*π, θ, π*). The corresponding exchange matrix is

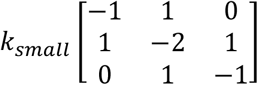

Rocking motions involve 6 symmetrically oriented sites corresponding to 6 distinct orientations of the H_2_O molecule, which approximately maintains its coordination within hydrogen bonding environment.[25] The explicit Euler angles of these sites are given by (0,0,0), (*±φ*, 2*π*/3 , 2*π ∓ φ*), (*π ± φ*, 2*π*/3 , *π ∓ φ*), and (*π*/2 , *π*/2 , *π*/2). The exchange matrix allows for a direct exchange between pairs of sites sharing the same orientation for one of the O*−*H bonds:

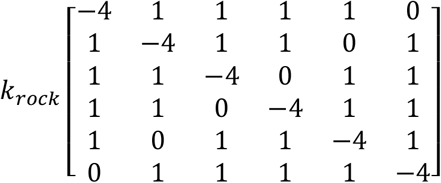

When a motional model calls for two types of motions occurring simultaneously, the overall motion is described through the nested frames with either the large-angle jumps or small-angle fluctuations frame being embedded in the rocking motion frame. A combined one-frame motion is then set up according to the procedure outlined in Vold and Hoatson.[20]

## Results and Discussions

### Overview of main experimental data and general model considerations

The main experimental data consists of ^17^O CT spectra, as well as ^17^O CT *T*_1_ and *T*_1_*ρ* relaxation times collected at 18.8 T and in the 240 to 180 K temperature range under static conditions (Figures 2-4). *T*_2_ times under static conditions are prone to proton decoupling imperfections and are not included in the analysis. We also discuss additional insights obtained from *T*_1_ and *T*_2_ relaxation times data under MAS conditions at 18.8 T and selected temperatures between 320 and 240 K with a spinning rate of 25 kHz.

**Figure 2.**
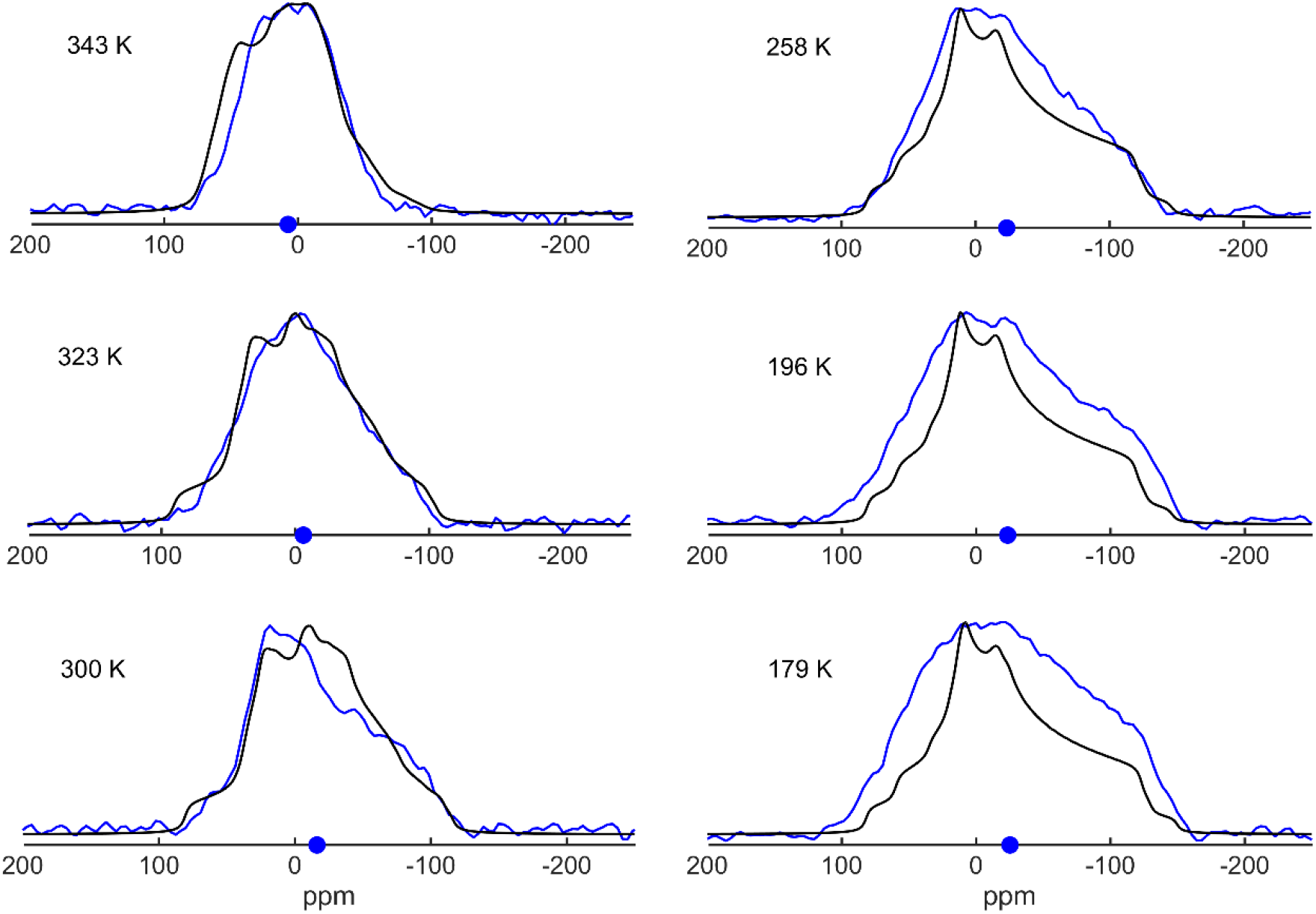
^17^O solid-state NMR line shapes of COM, collected under static conditions at 18.8 T (blue lines), overlaid after normalization with simulations according to the models described in the text (black lines).

**Figure 3.**
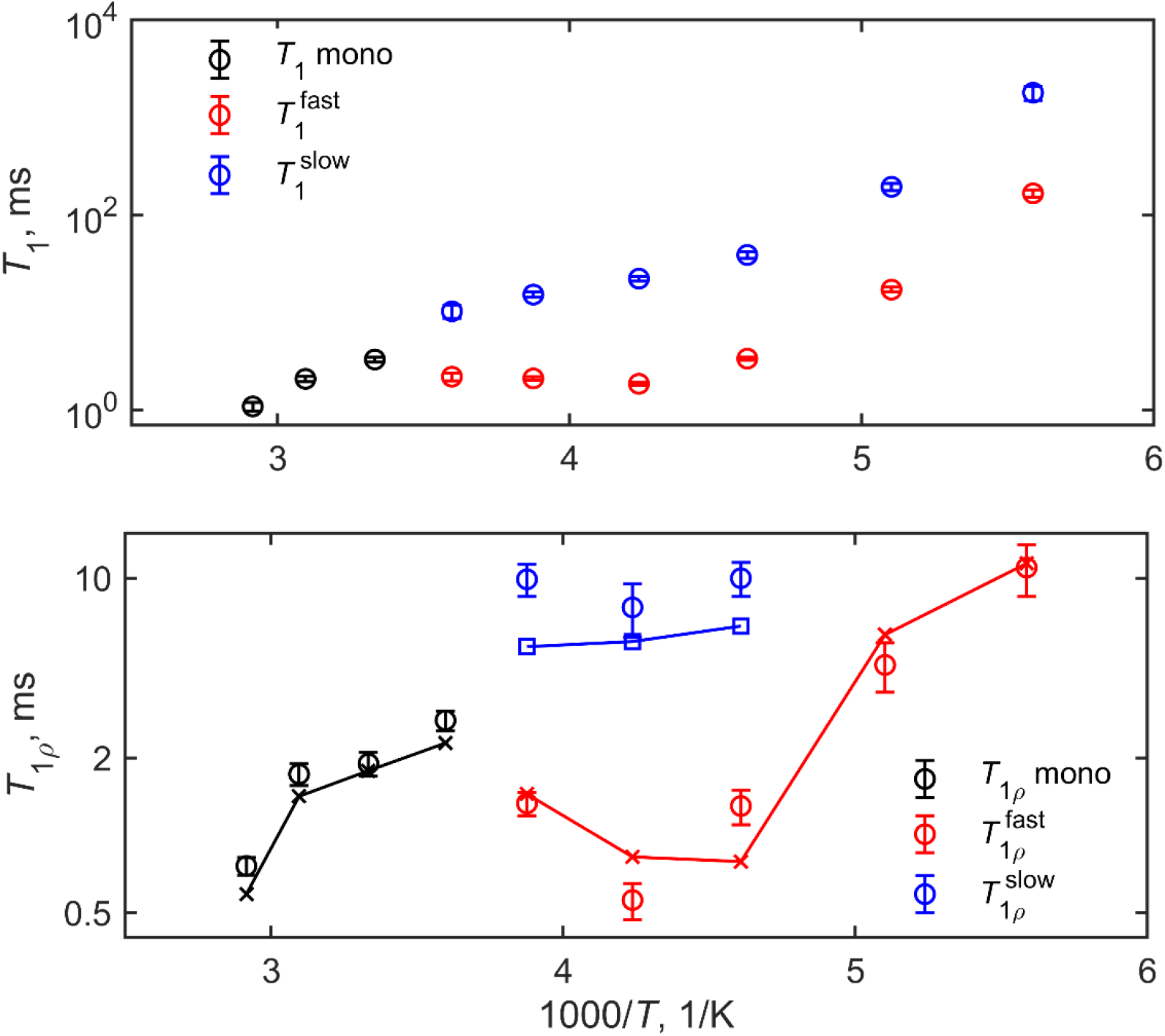
^17^O CT *T*_1_ times and *T*_1*ρ*_ times versus 1000/*T* collected under static conditions at 18.8 T. Integration over the entire powder pattern was performed to construct magnetization decay curves, which were fitted to mono-exponential decay/build-up above 300 K and the bi-exponential decay/build-up (Eq. 1) below 300 K. Corresponding simulations for *T*_1_*ρ* times using the model described in the text are shown by crosses connected by solid lines.

The ^17^O static solid-state NMR line shapes remain relatively similar between 280 to 180 K, but display a marked progressive narrowing in the 280 to 340 K range (Figure 2 and Figure S1A). This indicates that at high temperatures during the transition to the HT phase there is also an onset of large amplitude motions on the time scale of the quadrupolar coupling constant, which is about 6-7 MHz for both sites.[9] From the line shapes alone, the details of the dynamics is difficult to discern in the LT phase. However, the temperature dependence of the relaxation times indicate the presence of fluctuations, which must originate from small-angle motions within the constraints of the lattice. As a side point, temperature-dependent changes to the isotropic chemical shifts are observed, with significant changes evident upon the onset of the HT form. The isotropic chemical shifts are shown as blue dots in Figure 2 and plotted in Figure S1B.

It is interesting to observe the onset of non-exponentiality of the decay/build-up curves at temperatures below 280 K for *T*_1_ times and below 260 K for *T*_1_*ρ* times (Figure 4). Qualitatively, the non-exponentiality can originate either from differences in the dynamics in the two sites (W_2_ and W_1_), from the intrinsic non-exponentiality due to the differences in the relaxation rates for different crystallite orientations, or from the combination of the two factors. The large-angle motions in the HT form effectively average crystallite-dependence of the relaxation rates. The lack of non-exponentiality at high temperatures also suggests no significant differences between the dynamics of the two sites in the HT form, as suggested by the structural data. [5, 26]An alternative explanation is that the W_2_ and W_1_ sites still exhibit very different dynamics in the HT form, but due to fast exchange between water molecules, the relaxation is exponential; however, this explanation contradicts structural constraints. In the LT form, it is conceivable that the two sites experience a somewhat different extent of small-angle fluctuations due to differences in the local environment, although from the structural data it is not clear which of the sites is expected to be more mobile. In the discussion below, we label the less mobile of the two sites as “A” and more mobile as “B”.

**Figure 4.**
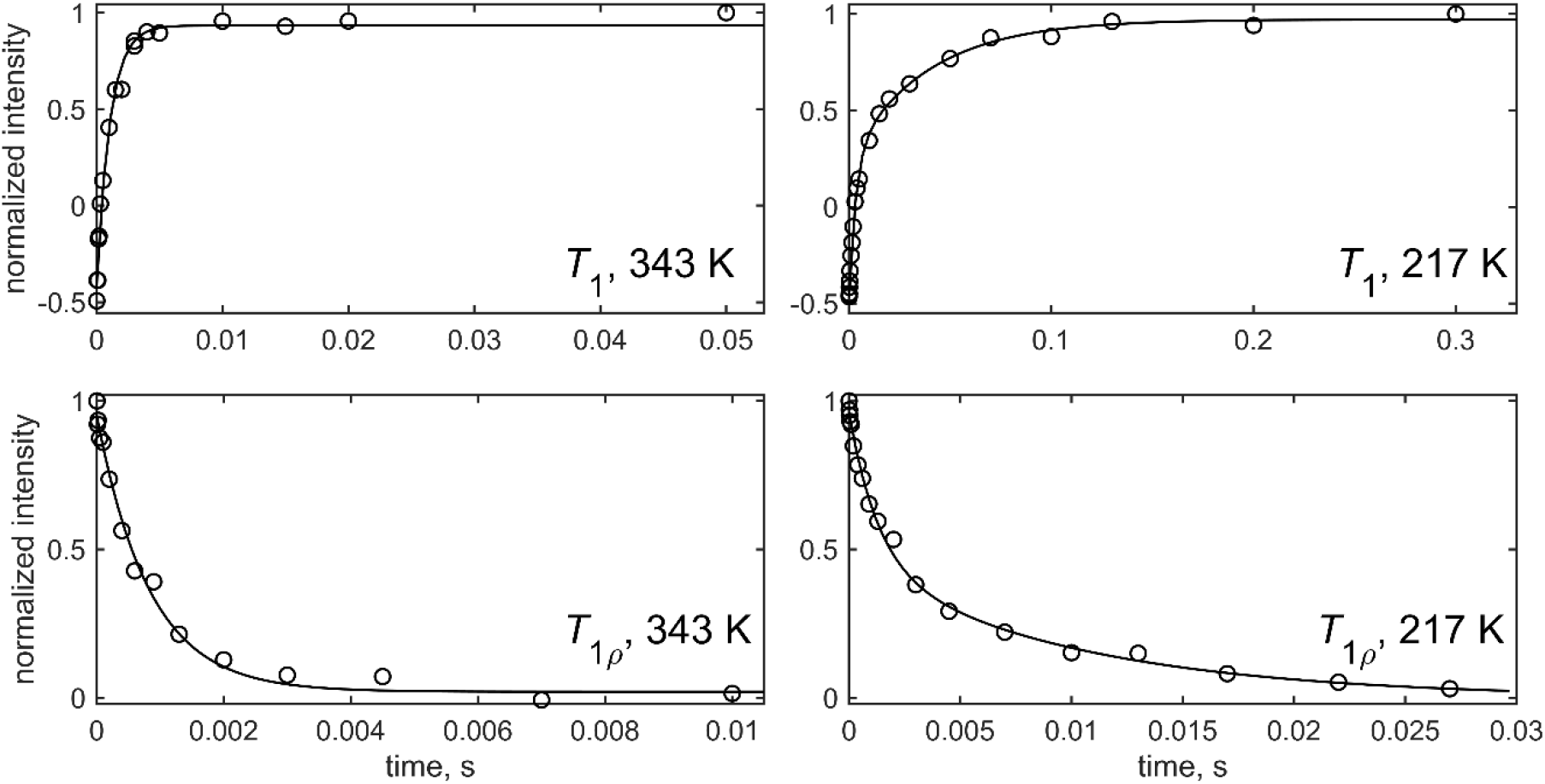
Examples of normalized magnetization build-up *T*_1_ or decay *T*_1_*ρ* curves at 343 and 217 K, showing the fits (solid lines) to either the mono-exponential function for the HT form at 343 K or the bi-exponential function for the LT form at 217 K. The intensities were obtained from integration of the entire powder patterns. The data were collected at 18.8 T under static conditions.

The differences in the dynamics of the two sites was also demonstrated in the *T*_2_ relaxation times measurements under MAS conditions by Goldberga et al.[9]

We model the dynamics in both forms of COM with the simplest model consistent with all experimental data (Figure 5) – 2-site large-angle jumps for the HT form (above 340 K) and small-angle fluctuations modeled as 3-site jumps for the LT form (below 280 K). The rate of jumps in the LT form can vary between the two sites. At the two intermediate temperatures of 323 and 300 K, the data suggests a combination of large- and small-angle jumps/fluctuations, which we interpret as a possible indication that one of the sites (labeled B) retains the features of the HT form, while the A-site begins to rigidify. For the large-angle fluctuations in the HT form we assume that in any given instant, the predominant motion is given by the distortions of one of the H-bonds in each water molecule, while for the second one the position is kept constant (tethered). As we will see below, our *T*_2_ data obtained under MAS conditions also points to an additional very slow collective mode of motions, which we denote as “rocking”. Interestingly, these types of rocking motions were also seen for protein crystals using solid-state NMR relaxation measurements.[27]

**Figure 5.**
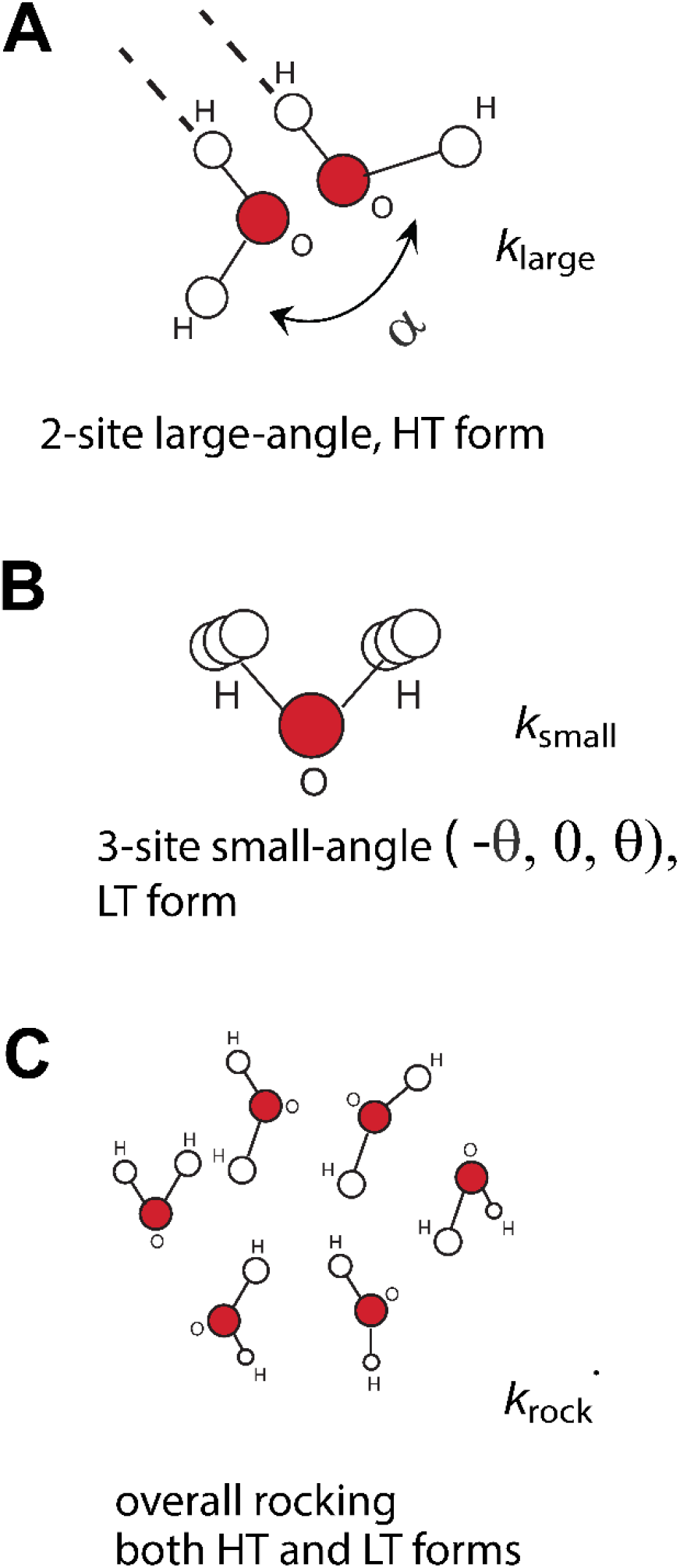
Motional models of the water molecules in COM: A) large-angle 2-site jumps with one of the hydrogen-bonds tethered, with the amplitude *α* and rate constant 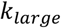 , predominant mode in the HT form. B) Small-angle fluctuations with the amplitude *θ* and rate constant 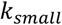 , dominant in the LT form. C) Overall collective very slow motions of the crystal with the rate constant 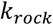, modeled as 6-fold tetrahedral reorientations.

### Details of model development and determination of motional model parameters

The EFG and CSA tensors for the two water sites were determined previously by Goldberga et al.[9], starting from MQMAS methods, and then performing a multiple field fit of both MAS and static data, at the set point range of 298-273 K. We use these values, listed in the *Material and Methods* section, as our main fits. The *C*_q_ values are around 6.3-6.5 MHz for both sites.

Simulated line shapes are constructed as the sum of the two underlying sites with equal populations. We first assume the fast motional limit in which the rate constant follows the constraint *k>>C*_q_, and determine the value of the 2-site jump amplitude consistent with the line shape at high temperature. The amplitude corresponds to 100*°* at 343 K, decreases to about 75*°* at 320-300 K, and to less than 15*°* at 280 K. *T*_1_ times allow for the determination of the values of the rate constants, and *T*_1_*ρ* measurements further constrain the model and the amplitudes of fluctuations.

At 343 K, the HT form is fully developed, and the 2-site large-amplitude jumps with the same rate constant for both W_1_ and W_2_ sites can explain all three measurements: static ^17^O line shapes, *T*_1_ and *T*_1_*ρ*. With the large-amplitude jumps of 100*°* the rate constant 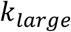 is 2 × 10^7^ s^-1^.

At low temperatures below 280 K, small-angle fluctuations with different rates for the two sites can fit the data. The need for two separate rate constants for the W_1_ and W_2_ sites is demonstrated by the non-exponentiality of the *T*_1_ magnetization build-up curves (Figure 4). The combination of *T*_1_ and *T*_1_*ρ* data constraints the amplitude of motions, as the ratio *T*_1_/*T*_1_*ρ* at each temperature in the slow motional regime is dependent on the amplitude.[12]

In order to match the fast and slow components of the *T*_1_ relaxation, the B site requires that the value of small-angle fluctuations amplitude *θ* be 8*°* or less between 280 and 200 K (Figure S2). The A site allows for more flexibility in fitting *T*_1_ times in this range, with *θ* in the range of ca. 8-20*°*. However, further *T*_1_*ρ* constraints impose *θ*=8*°* for this site in the 260 to 200 K range. Below 200 K, the *T*_1_*ρ* relaxation decay curves are not consistent with the 8*°* amplitude model and instead require the values of *θ*=4*°* at 196 K and *θ*=2*°* at 179 K. The examples of *T*_1_*ρ* magnetization decay are shown in Figure 4, and Figure S3 show examples of how the values of amplitudes were constrained by the *T*_1_*ρ* decays at low temperatures. With this approach, the fitted values of *k*_*small*_ display roughly an Arrhenius temperature dependence (Figure S4) with the activation energy values of 14 kJ/mol for B site and 10 kJ/mol for the A site.

For the intermediate temperatures of 323 and 300 K, it is not possible to choose a single mode of large-angle jumps to fit all the data. The line shapes and *T*_1_ curves can be reconciled with the single mode of large-angle jumps, but not the *T*_1_*ρ* curves, which display much slower decays if only the motion with *α*?=75-50*°* (which fits the line shapes and *T*_1_ times combination) is invoked. A better consistency between all three types of measurements at these temperatures is attained if we assume that the B site retains the large-angle mode with *α*=75*°*, but the A site begins to rigidify earlier and exhibits only the small-angle fluctuations with *θ*=15*°*(Figure S4). The differences in the dynamics of two-sites can be in principle obtained from molecular dynamics (MD) simulations, but care must be taken to use approaches that can quantitatively probe slower time scales.

**Table 1.** Summary of motional modes and fitted parameters for COM in the 343 to 180 K range based on ^17^O CT line shapes, *T*_1_ and *T*_1_*ρ* times at 18.8 T under static conditions.

| Temperature range | A, less mobile site | B, more mobile site |
| --- | --- | --- |
| above 343 K, HT form | Large-angle jumps with $\alpha=75$-$100^\circ$<br>$k_{large} > 2 \times 10^7 \text{ s}^{-1}$ | |
| 325-300 K | Small-angle jumps $\theta = 15^\circ$,<br>$k_{small}$ range $8-3 \times 10^7 \text{ s}^{-1}$ | Large-angle jumps with $\alpha = 75^\circ$,<br>$k_{large} = 1.1 - 0.7 \times 10^7 \text{ s}^{-1}$ |
| 280 K-220, LT form | Small-angle jumps $\theta = 8^\circ$,<br>$k_{small}$ range $2 \times 10^9 - 2 \times 10^7 \text{ s}^{-1}$<br>rocking motions, $k_{rock} = 50-100 \text{ s}^{-1}$ | |
| 220- 180 K, LT form | Small-angle jumps $\theta = 4-2^\circ$,<br>$k_{small}$ range $2 \times 10^8 - 6 \cdot 10^6 \text{ s}^{-1}$ | |

Additional *T*_1_ times were measured under 25 kHz MAS conditions (Figure S6) for a limited range of temperatures due to probe limitations (330 to 235 K). We also measured *T*_2_ times at 325, 258, and 235 K with the application of SPINAL-64 ^1^H decoupling during the relaxation delay period. (Figures 6 and S7). *T*_1_*ρ* times under MAS conditions are more complicated to interpret due to MAS-driven interconversions of coherences and are not included in the present work. However, *T*_1_ relaxation times under MAS are consistent with results under non-spinning conditions. (Figure S6).

**Figure 6.**
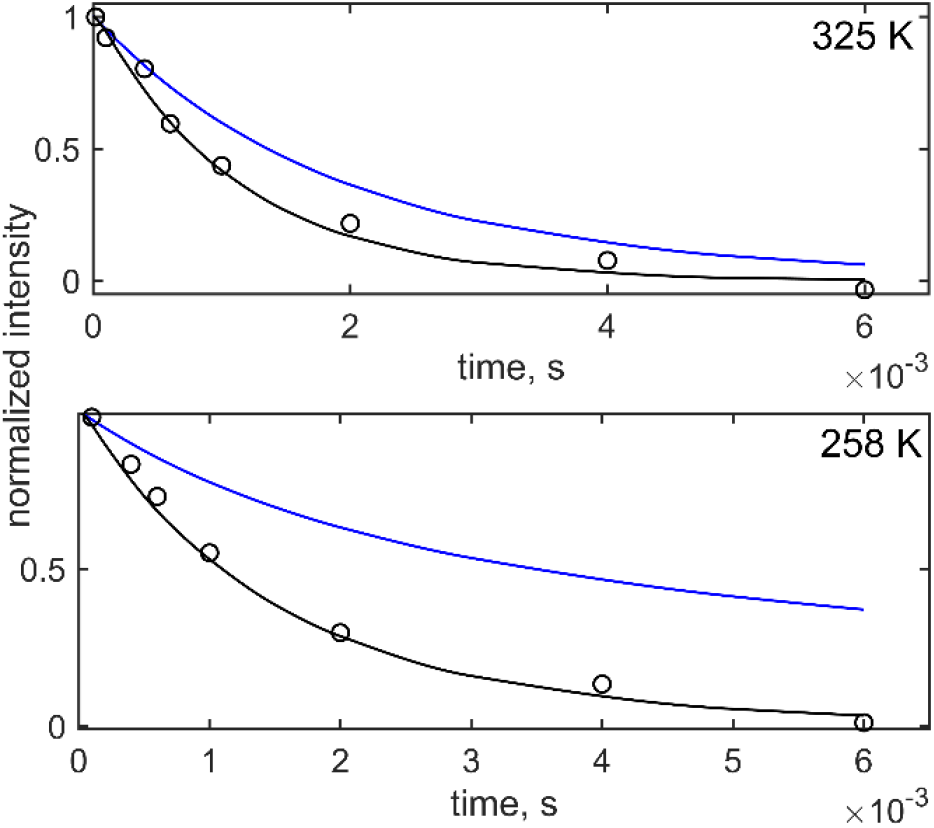
Normalized *T*_2_ relaxation decay curves at 325 and 258 K, obtained at 18.8 T and 25 kHz MAS rate. Experimental intensities corresponding to the integration of the entire central band (black circles) are overlaid with simulations according to the model described in the text and either with the inclusions of the symmetric “rocking” motions with *k*_*rock*_=100 s^-1^ (black lines) or without this motion (blue lines).

However, comparing experimental *T*_2_ magnetization decay curves with those simulated according to the model and parameters derived on the basis of data under static conditions shows a marked qualitative discrepances: at 325 K, the simulated curve decays are slower than the experiment and at lower temperatures simulated data displays a much slower long component not observed in the experiment (Figures 6 and S7). The absence of the slow component signifies presence of a symmetric motional mode, referred to as “collective rocking motions” in literature.[27] As a modeling approximation for this, we invoke an approach using 6-fold symmetrical tetrahedral reorientations.[25] A rate constant 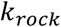 on the order of about 50*−*100 s^-1^ is sufficient to render qualitative agreement with the experimental data (Figures 6 and S7) at all temperatures. The inclusion of this mode does not affect either the simulated static line shapes or *T*_1_ values. It also does not affect static *T*_1_*ρ* simulations above 220 K when the value of 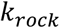 is between 100 to 50 s^-1^. However, for two low temperatures of 196 and 179 K, *T*_1_*ρ* simulations are affected and do not correspond to the experiment. We thus conclude that the collective rocking motions mode likely freezes below about 220-200 K. Interestingly, this is close to the temperature known to cause the onset of freezing of the hydrogen bonding network in various amorphous water-containing systems and ice.[28]

Relaxation data under MAS in the 280-240 K range shows a tendency for differential relaxation times in each of the two sites (Figure S8). However, the resolution is not sufficient to obtain individual components for each site. Rather, as in the case of the relaxation decay curves under static conditions, the integrated spectral intensity was fitted to the bi-exponential fitting function. A similar approach was undertaken by Goldberga et al.[9] for *T*_2_ data analysis under 35 kHz MAS at the set point temperature of 273 K. Our *T*_2_ data under 25 kHz MAS also shows a tendency for individual components for each of the sites, but within the quality of the data, the integrated powder pattern can be fitted with the single exponent, and a double-exponential fit would be an overfitting. The values of the single-exponential *T*_2_ times are 1.29±0.16 ms, 1.67±0.14 ms, and 1.49±0.16 ms for 325, 258, and 236 K respectively.

Additional subtle nuances of motions can be obtained by careful comparison of the simulated line shapes with the experimental data at low temperatures (Supporting Information SI-1 and Figure S9): the experimental spectra are wider and have shapes corresponding to a smaller quadrupolar asymmetry parameter. This was already noted by Goldberga et al.,[9] and the effect is more pronounced in our work, as our lowest temperature is 179 K, accounting for calibrations using lead nitrate. The site-specific MQMAS values were obtained at a much higher “set point” temperatures of 273-298 K and the discrepancies between the simulations with these parameters and the low temperature line shapes indicate the freezing of additional motional modes, although the rigid values of the tensors as calculated by the DFT approach[10] are still not attained at 180 K.

An alternative model for the HT form can follow from the structural data using the four-site model[5], shown in Figure 1B. While the line shape and *T*_1_ relaxation time at 343 K can be fit adequately with this model, there is a significant discrepancy with the *T*_1*ρ*_ relaxation time data (SI-2 and Figure S10).

**In conclusion**, the combination of ^17^O NMR line shape analysis, along with *T*_1_, *T*_2_ and *T*_1_*ρ* relaxation times over the wide temperature range allowed for a detailed examination of water dynamics in COM, complementing prior structural studies of this important mineral. Line shape and *T*_1_ data provided initial constraints on the types of motion (large angle-jumps in the HT forms and small-angle fluctuations in the LT form), while the *T*_1_/*T*_1_*ρ* ratio restricted the amplitudes of motions and provided precise constraints on the rate constants. This ratio also constrained the behavior in the intermediate temperature region between 325 to 300 K, requiring large-angle motions to freeze in one of the sites, while large-angle jumps, albeit with a smaller amplitude, persist in the second site. *T*_2_ data indicated the existence of very slow collective “rocking” motions down to about 200 K, while the *T*_1_*ρ* decay curves defined the low temperature limit at which these rocking motions are no longer consistent with the data.

## Acknowledgements

This work was supported by National Institutes of Health grant 1R15GM157635-01A1 to L.V. All solid-state NMR experiments were performed at the National High Magnetic Field Laboratory, which is supported by NSF Cooperative Agreement NSF/DMR-2128556, the State of Florida, and the U.S. Department of Energy.

We are grateful to Danielle Laurencin and Ieva Goldberga for providing the COM sample, as well as helpful discussions of the work.

