## Supporting Information for "Dynamics of calcium oxalate monohydrate in high and low temperature phases using ^17^O solid-state NMR"

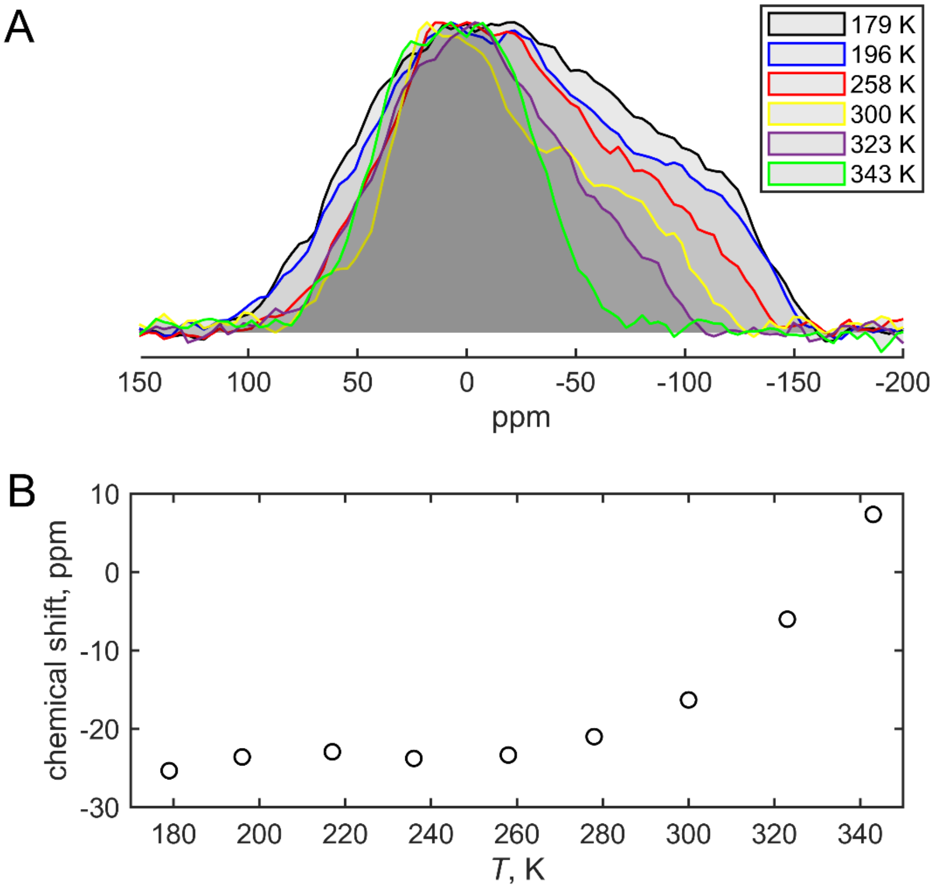


**Figure S1.** A) Overlay of normalized experimental spectra at selected temperatures: the same spectra as in Figure 2 of the main text are shown. B) Temperature dependence of the "center of mass" weighted isotropic chemical shifts from the line shape fits. At 300 K this weighted chemical shift value is -16 ppm, which is within the error limits to the value of calculated from the data of Goldberga et al. [1]

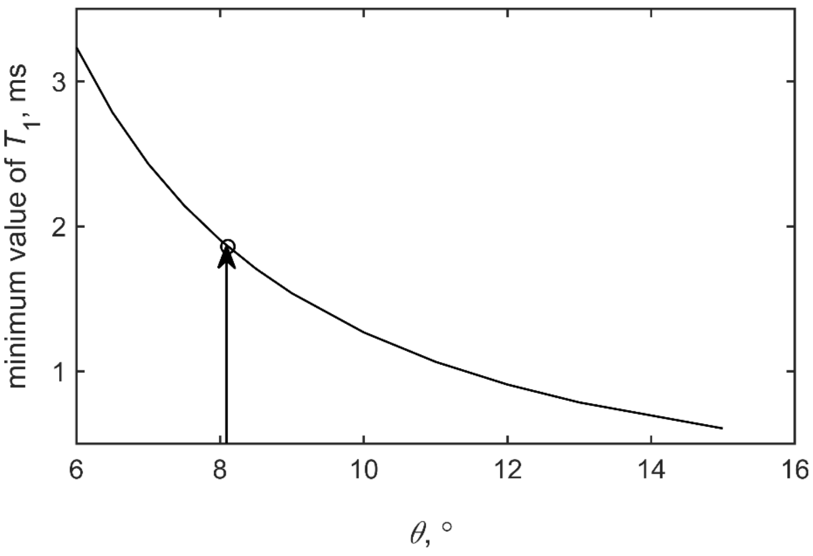


**Figure S2.** Minimum value of the longitudinal relaxation time, $T_1$, as a function of the amplitude of small-angle fluctuations for the more mobile B site. The simulation parameters are given in the main text. For all angles, $T_1$ minimum is reached for $k_{\text{small}}$ = $10^9$ $s^{-1}$. A circle and a vertical line indicate the value of $T_1$ and the corresponding amplitude of small-angle fluctuations for the minimum value of the fast component of $T_1$ build-up curves obtained in the experiment.

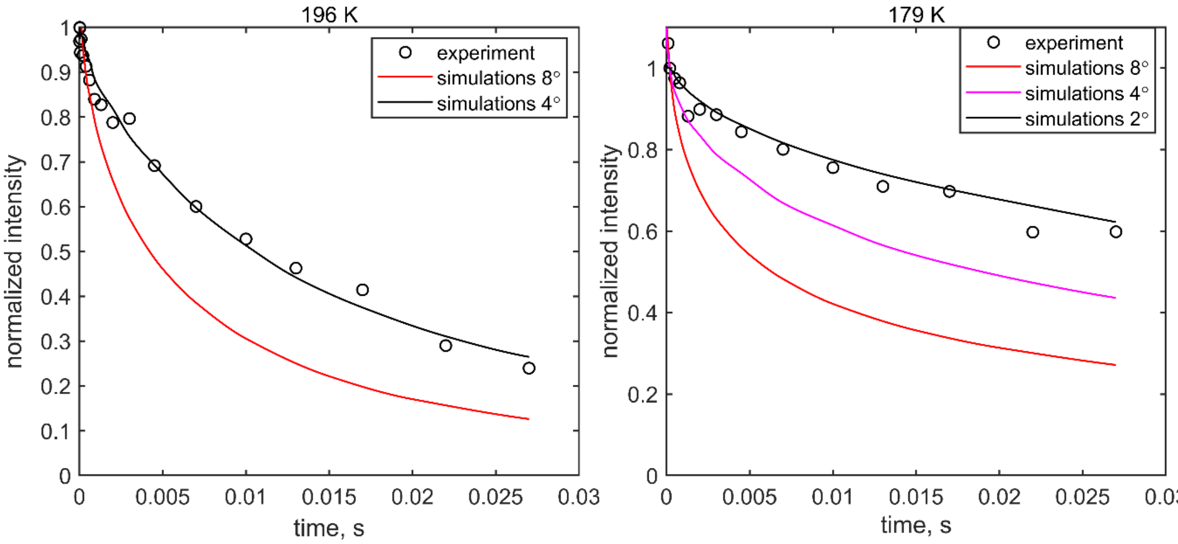


**Figure S3.** Experimental (circles) and simulated relaxation decay curves for $T_{1\rho}$ at 196 K (left panel) and 179 K (right panel). Simulated curves are calculated using the model of small-angle fluctuations with the amplitude of either $\theta$= 8° (red line) or $\theta$= 4° (black line) for 196 K and either $\theta$= 8° (red line), $\theta$= 4° (magenta line), or $\theta$= 2° (black line) for 179 K. The values of $k_{small}$ used in these simulations were taken from the fits to the experimental $T_1$ values, as elaborated in the main text.

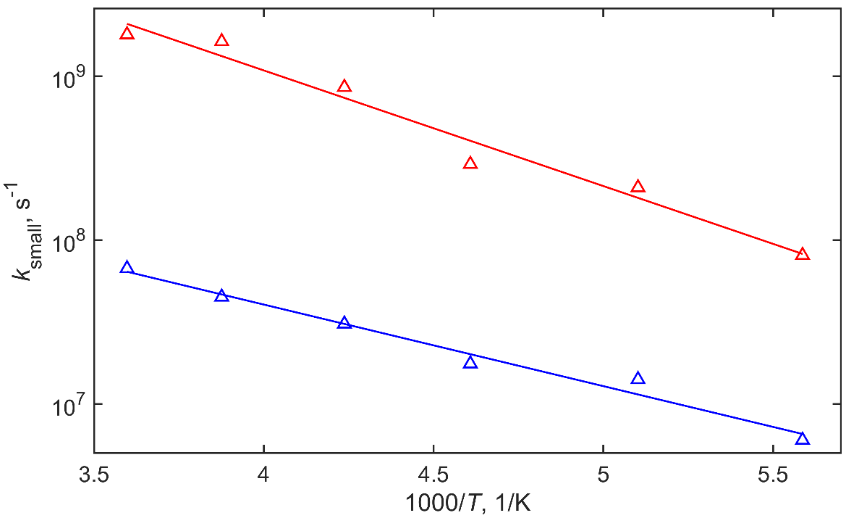


**Figure S4.** An Arrhenius plot of the fitted values of $k_{small}$ for the temperatures below 280 K. Fitted values at individual temperatures (triangles) are obtained by fitting the slow (blue) and fast (red) relaxing components of $T_1$ build-up curve. Straight lines show the Arrhenius fits with activation energies 9.5±0.7 kJ/mol and 13.5±1.2 kJ/mol, respectively.

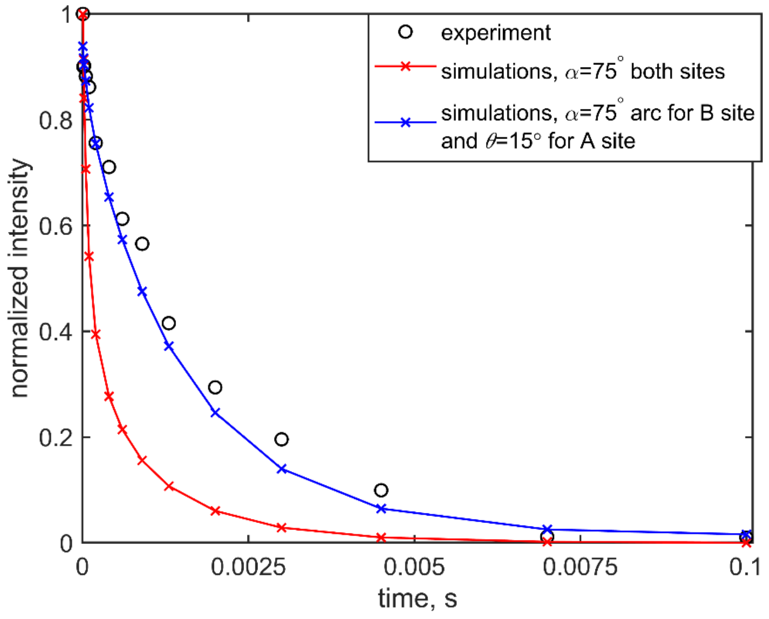


**Figure S5.** Experimental (black circles) and simulated $T_{1\rho}$ relaxation decay curves at 323 K. Simulated curves are calculated for the model of large-angle jumps with $\alpha$ = 75° in both sites (red line and crosses) or large-angle jumps with $\alpha$ = 75° in B site and small-angle fluctuations with $\theta$ = 15° in A site (blue line and crosses). Both types of simulations also include rocking motions with $k_{rock}$ = 100 $s^{-1}$. The rate constants of large-angle jumps and small-angle fluctuations were obtained from fits to experimental $T_1$ values, as explained in the main text.

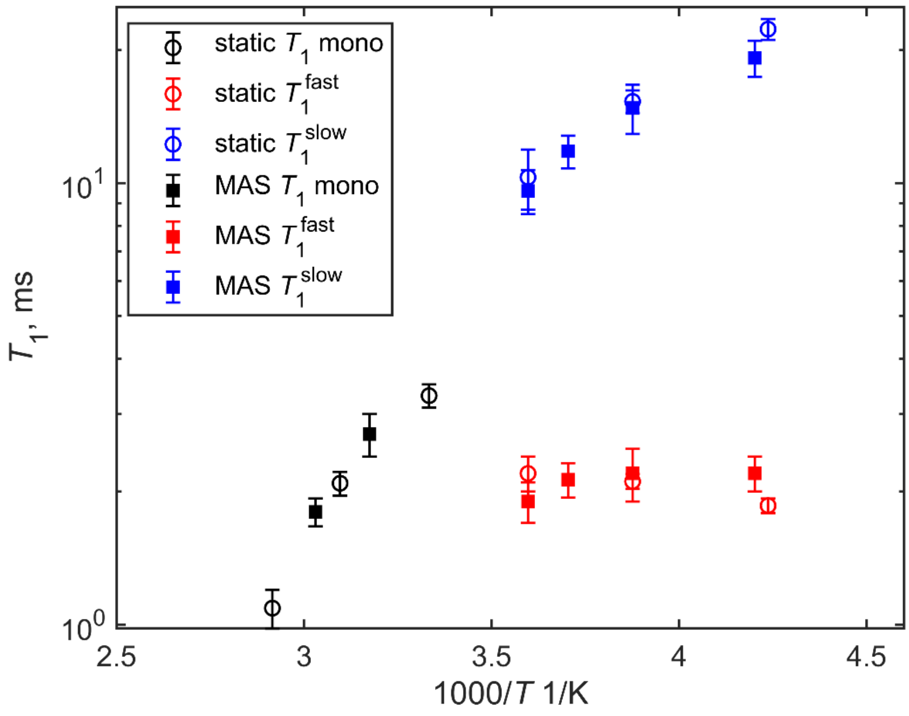


**Figure S6.** Overlay of $T_1$ times versus 1000/$T$ obtained either under static (circles) or MAS (filled squares) conditions at 18.8 T. The MAS rate was 25 kHz.

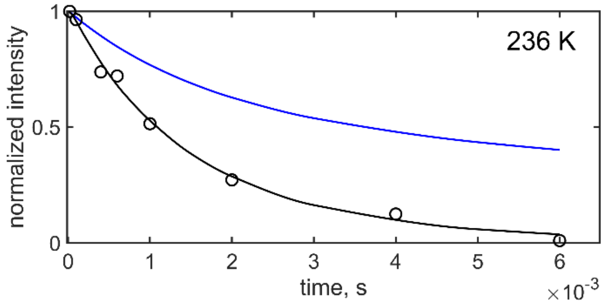


**Figure S7.** Normalized $T_2$ relaxation decay curve at 236 K obtained at 18.8 T and 25 kHz MAS rate. Experimental intensities corresponding to the integration of the entire central band (black circles) are overlaid with simulations according to the model described in the text and either with the inclusions of the symmetric “rocking” motions with $k_{rock}$=100 s$^{-1}$ (black line) or without this motion (blue line).

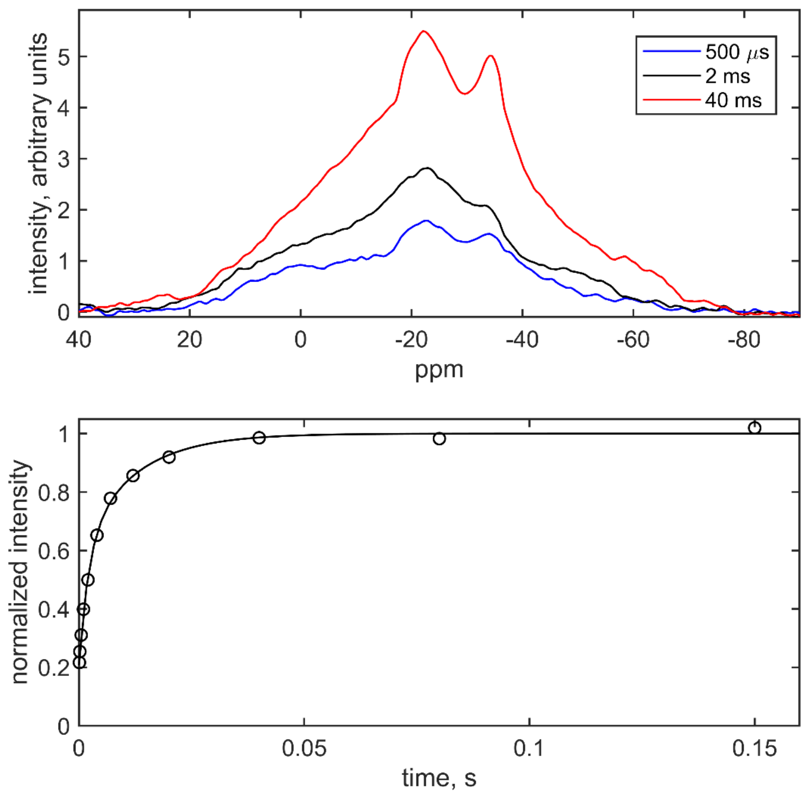


**Figure S8.** Details of $T_1$ relaxation data collected at 25 kHz MAS rate at 18.8 T. Top panel) The tendency for differential relaxation in the low temperature $T_1$ data, demonstrated in the partially relaxed spectra at 258 K. The relaxation delay times are shown directly on the panel. In this case 0.1 kHz line broadening was applied to enhance the resolution. MAS has a negligible effect on the relaxation rates (Figure S5), as was also supported by the simulations. Bottom panel) Due to low resolution, the two individual components were not obtained from these spectra, but rather from the double-exponential fit of magnetization build-up curve, shown for 258 K. The line represents the fit to the double-exponential function, with the fitted $T_1$ values of 15.3±0.09 ms and 2.11±0.08 ms.

#### SI-1. Additional analysis of EFG tensors from low temperature $^{17}O$ NMR line shapes.

As noted in the main text, the experimental spectra at low temperatures are wider and have shapes corresponding to a smaller quadrupolar asymmetry parameter when using the values reported by Goldberga et al.[1] Instead, if an equal increase in $C_q$ is applied to both sites, the fit can be obtained with the following values for the quadrupolar tensor parameters $C_q$ = 7.6 MHz, $\eta_q$ = 0.39, for the site $W_1$ and $C_q$ = 7.5 MHz, $\eta_q$ = 0.58 for the site $W_2$, as shown Figure S9A. In this approach, the parameters of the CSA tensors were kept unchanged, as shown in Goldberga et al.[1] We have chosen not to employ these corrections in our main fits because a) they still do not reach the rigid values calculated on the basis of the DFT approach[2] and b) we cannot obtain true site-specific information without the assumption of the uniform scaling. The DFT values are $C_q$ = 7.6 MHz, $\eta_q$ = 0.98 for site $W_1$ and $C_q$ = 8.7 MHz, $\eta_q$ = 0.81 for site $W_2$. We investigated the effect of changes in the tensor parameters on the relaxation rates, and while there are obviously quantitative changes in the fitted values of the rate constants, there is no trend for qualitative differences. For example, one still needs to employ different $k_{small}$ rates for both sites, the amplitudes of small-angle fluctuations still need to be reduced below 220 K, and the removal of the long tail in MAS $T_2$ decay curves still cannot be achieved without invoking a symmetrical motion. The additional narrowing of the line shapes to match the experiment at high temperatures when these modified tensors are used can be achieved by invoking an additional anisotropic mode (Figure S9B, shown for 278 K), for example a restricted motion inside a potential well or the so-called "arc motion"[3] with the amplitude of about $\alpha = 30°$ and varying rate constants $k_{arc}$ (Figure S8C). The arc itself is represented by several sites with values of rotation angle incremented in $\Delta\alpha = 5°$ steps. The exchange rate between nearest-neighbor sites is given by $k_{arc}(\alpha/\Delta\alpha)^2$.

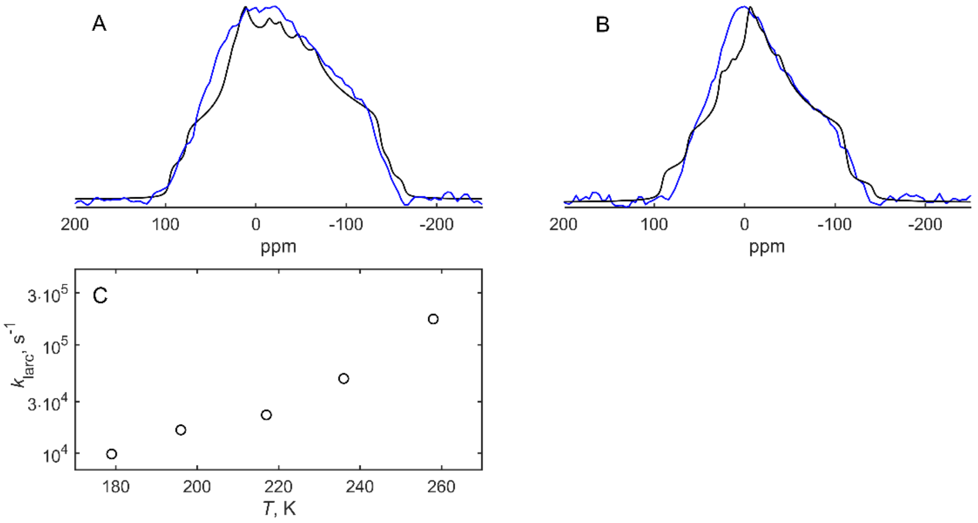


**Figure S9.** Experimental line shapes at A) 179 K and B) 278 K overlaid with simulations using the modified EFG tensor parameters, listed in SI-1. Simulations (black lines) at 278 K include small-angle fluctuations with 8° amplitude as well as the anisotropic "arc" mode of motion with the amplitude of 30° and the rate constant $k_{arc}$ of $3.1\cdot 10^5$ s$^{-1}$. Simulations at 179 K include only the small-angle fluctuations with $\theta = 2°$ amplitude. C) Temperature dependence of $k_{arc}$ based on fits of the experimental line shapes with the use of the modified tensors.

### SI-2. Alternative HT form model

The structural model for the HT form suggested by Shepelenko et al.[4] includes four orientations for a water molecule in a single site related by the mirror reflection and a 180° rotation around the axis perpendicular to the mirror plane. In terms of $^{17}O$ quadrupolar tensor PAS, these four orientations are given by the following Euler angles: (137, 108, -5), (137, 72, 5), (135, 81, 100), and (135, 99, 80). The alternative dynamics 4-site model consists of jumps between these four orientations. This model was used with the EFG and CSA tensor parameters separately for the $W_1$ and $W_2$ sites, and the obtained line shapes and relaxation curves were then averaged.

The experimental value of $T_1$ time of 1.09 ms at 343 K can be fitted with the rate constant $k_{HT} = 1.0\cdot 10^7$ s$^{-1}$. The simulated line shape obtained with these parameters is shown in Figure S10 and is reasonable although somewhat worse than the main fit with the 2-site model. However, with this rate constant the simulated $T_{1\rho}$ relaxation time is 0.072 ms which is a factor of 10 smaller than the experimental value of 0.76 ms.

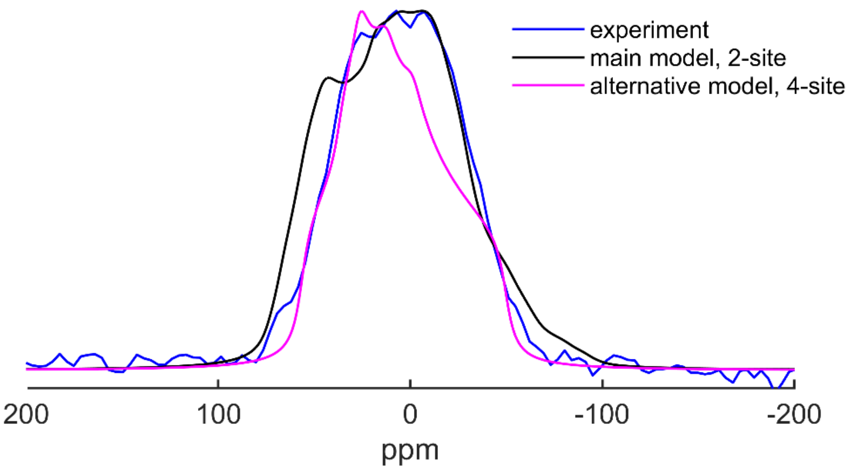


**Figure S10.** Comparison of normalized experimental $^{17}$O CT line shape (blue line) at 343 K with the simulations according to the 4-site alternative HT model using the rate constant of $k_{\mathrm{HT}} = 1.0 \cdot 10^{7}$ s$^{-1}$, obtained from the fit of the $T_1$ relaxation time (purple), as well as the 2-site large-angle jumps model of the main text (black line).
